# Feather degradation in *Stenotrophomonas maltophilia* relies on conversion and internalization of keratin monomer

**DOI:** 10.1101/2024.03.22.586301

**Authors:** ZhangJun Cao, XiaoXiao Song, Kai Xue, Wei Zhang, YunLong Zhang, Ting Chen, XingQun Zhang

## Abstract

Feather keratin is the most abundant nitrogen source waste in nature. This insoluble material cannot be directly utilized as nutrition by most organisms, especially animals and plants. While feathers are naturally decomposed by some microorganisms through keratinase-based degradation which remains mechanistically not fully understood. In this study, we find that when feathers serve as the only nutrient source for *Stenotrophomonas maltophilia* culture, keratin monomer of about 10 kDa is present in the medium as a predominant hydrolyzed product. We further show that keratin monomers bind to cells and in turn enter cells via an undetermined pathway. The cell entry of keratin monomer elicits keratinase activity to promote the forward reaction in keratin hydrolysis. This study highlights the importance of keratin monomer production as the first step in feather biodegradation, in which the insoluble feather is converted into soluble intermediate to facilitate its internalization and complete hydrolysis.

**Importance:** A large amount of feathers have been produced in poultry breeding, which could cause environmental pollution if not processed in time. On the other hand, amino acids degraded from feathers can be used in many fields, such as feed, fertilizer, daily chemicals and other fields. Biodegradable feathers have the advantages of low energy consumption, mild conditions and no destruction of the amino acids obtained from degradation. We previously isolated and identified a highly efficient feather-degrading bacterium, Stenotrophomonas maltophilia DHHJ, which can grow well on the medium with feathers as the only nutrient and completely degrade feathers. We know, feather particles are much larger than bacterial cells. In what form and how they are absorbed into cells by bacteria are interesting and critical questions for feather degradation. In our research, feathers had been first degraded extracellularly by basically expressed keratinase into keratin monomers. The keratin monomers bound to cells and enter across the membrane and can induce further expression of keratinase. The understanding of keratin monomers provides new clues for the study of feather degradation mechanisms.

## Introduction

A huge amount of keratin waste is generated annually as a by-product from many different sources such as poultry farms, slaughterhouses, and textile industry. In 2019 alone, more than 4.7 million tons of chicken feathers were produced from the global poultry industry(1). Uncurbed dispose and accumulation of feather waste have posed a worldwide challenge for our environment (2). On the other hand, feather waste is a protein-rich material with great value for recycling, because feathers can be converted to value-added products in many industrial fields such as cosmetics, organic fertilizer and livestock feed. Physical (high temperature, explosion and microwaving) (3) and chemical treatments (hydrogen peroxide, acid or alkali) (4, 5) or their combination (6) have been employed as the conventional methods for degradation and decomposition of feathers. However, these treatments, which normally require harsh processing conditions, are prone to cause poor recovery yield of essential amino acids in the product due to molecular destruction (7, 8). Additionally, high energy-consumption as well as environmental concerns are also hurdles for their commercial application (9).

In contrast to physical and chemical processes for feathers, biodegradation by microorganisms holds enormous promise for developing a green alternative, which has advantages, such as mild reaction conditions, efficient product recovery, environmental friendliness, and energy saving (10). Sustained efforts have led to the discovery of many keratinase-producing microorganisms as potent natural decomposers for these hard-degrading fibrous proteins (2). The most common kerotinolytic microorganisms include bacteria (most as gram-positive genera) and fungi, which can use keratin-rich substances as nutrition source for survival via a keratinase-dependent proteolytic process. The bacterium *Stenotrophomonas maltophilia* DHHJ strain, isolated in our laboratory (11), can grow well in the culture medium with feather powder as the sole nutrient source (carbon and nitrogen) by completely degrading the feather powder. Considering the large size of insoluble feather powders compared to bacterial cells, how do they enter bacterial cells to serve as nutrient molecules? How is the degradation enzyme system of bacterial cells activated? The answers to these questions are obviously the key to establish an efficient system for feather protein degradation.

The major component of avian feathers is β-keratin, an insoluble structural protein consisting of densely packed polypeptides highly cross-linked with disulfide bonds (12). Feather keratin was previously shown to be solubilized under certain conditions to yield a unit (named as keratin monomer) with a molecular weight of about 10 kDa (13). Although many feather keratin genes from *Gallus gallus* have been located on multiple chromosomes, each of these loci encodes a polypeptide of about 97 amino acid residues with very similar structure (14–16), closely resembling keratin monomer in size.

The special structure features confer the keratin’s resilience to the common proteases. Instead, keratinases produced in various microorganisms are proteolytic enzymes dedicated for digestion of keratins such as feathers, nails, hair, and wool (17). Extensive protein purification works have been done for various keratinases from different sources. Some purified keratolytic enzymes show the robust capability to break down keratins in vitro, while it is not surprising to see that keratinases are often less potent after separation from the crude extract (9, 18, 19), suggesting that the full function of keratinases likely rely on other factors in a system. A mechanism has recently been proposed that the complete feather degradation process may include two major steps: sulfitolysis and proteolysis (20, 21).

Here we intended to uncover the mechanism underlying keratin hydrolysis by *S. maltophilia*, a bacterium with desired capability in feather degradation (11). We found that keratin monomer will be generated at the initial stage of feather degradation and thereafter translocated into *S. maltophilia* cells for further degradation.

## Materials and Methods

### Bacteria and Culture Conditions

#### Stenotrophomonas maltophilia

DHHJ was grown in feather meal medium (feather meal 20 g/L, NaCl 5 g/L, K_2_HPO4 3 g/L, KH_2_PO4 4 g/L, pH 7.0-7.2), beef extract-peptone medium (beef extract 5 g/L, peptone 10 g/L, NaCl 10 g/L, pH 7.0-7.2) or M9 keratin medium (NH_4_Cl replaced with 0.4∼0.5 g/L keratin monomer, pH 7.0-7.2) incubated aerobically at 35 °C for 1–3 d. *E.coli* was grown in LB broth media for 14-16 h.

### Chemical preparation of feather keratin monomer

Feather keratin monomer (FKM) was prepared as described previously (Zhou et al. 2014). Briefly, 1 g of degreased feather was immersed in 10 ml of a reaction solution containing 0.125 mol/l Na_2_S_2_O_5_, 0.05 mol/l SDS, and 2.0 mol/l urea. The mixture was heated to 80 °C for 30 min. Then the solution was diluted with 40 ml of distilled water, filtered, and dialyzed against distilled water using a 12-14 kDa cutoff membrane for 2 h.

### Feather keratin monomer prepared in *E. coli*

The cDNA of feather keratin gene (Gene ID: 769269) was synthesized and cloned into pET-32(+) vectors to bacterially express a protein that contains N-terminal tandem protein tags: TrxA-6xHis-thrombin cleavage site-S tag (Fig. S1). After purification with Ni affinity chromatography, the recombinant keratin proteins were digested with thrombin protease to release S-tagged keratins that were further cleaned up with Ni column to eliminate the needless residues.

Bacterial total protein, membrane protein and intracellular protein extraction kits were purchased from Bestbio (Shanghai, China). Thrombin and enterokinase were from Yeasen (Shanghai, China). Beyo ECL Moon and anti-S Tag mouse monoclonal antibody were obtained from Beyotime Biotech Inc. (Shanghai, China). HRP and colloidal gold labeled rabbit anti-mouse IgGs were products from BBI Biotech (Shanghai) Co. Ltd (Shanghai, China).

### FITC (Fluorescein 5-isothiocyanate) protein labeling

FKM solution was dialyzed 3 times at 4 °C in a cross-linking reaction solution to reach pH 9.0. FITC solution was freshly prepared by dissolving in DMSO at a concentration of 1 mg/mL and protected from light. The labeling reaction was performed in the dark at 4 °C for 8 h, in which the FITC reagent was slowly added into the protein solution in a ratio of 100 μg (FITC):1mg (protein) by gently shaking. Then 5 mol/L NH_4_Cl was added to the abovementioned reaction mixture to a final concentration of 250 mmol/L, incubating at 4℃ for 2 h to terminate the reaction. The cross-linking mixture was dialyzed against PBS with buffer changes (more than 4 times) until the dialysate was clear.

### Cells incubation with keratin monomer

A 5-mL aliquot of *S. maltophilia* DHHJ culture in logarithmic growth phase was harvested and centrifuged at 5000 rpm for 5 min to collect bacterial cells. The cell pellets were washed twice with PBS, then resuspended with 5 mL of 50 mM Tris-HCl (pH 7.8) and 15 mL ddH_2_O, with addition of 5 mL FITC labeled keratin (fluorescence assay) or S-tag keratin (western blotting and immunocolloidal gold assays). These bacterial cells incubated at 35℃ or 4℃ were examined for fluorescence value using Hitachi F-4500 fluorescence spectrophotometer at 1 h, 2 h, and 3 h, respectively. The incubation was performed at 35℃ for 1 h prior to fluorescence microscope observation, western blotting, and immunocolloidal gold experiment. Unless specified otherwise, 400 ug/mL keratin was used in these reactions.

### Determination of fluorescence value

After incubation with FITC labeled keratin, cells were precipitated by centrifuging at 10,000 rpm for 2 min, followed by two washes with incubation buffer (IB, 10 mM Tris-HCl, pH 7.5). Then the cells were resuspended in equal volume of IB. The fluorescence value was measured with the Hybrid Multimode Microplate Reader (Synergy 2, BioTek Instruments, Santa Clara, USA) with the wavelength setting of 485 nm (excitation)/528 nm (emission).

### Western Blot analysis

After incubation with S-tag keratin, cells were washed twice with PBS, from which total, membrane, and intracellular proteins were separately extracted. The samples were resolved by SDS-PAGE (4–20% gradient gel) characterized by western blot analysis using HRP conjugated rabbit anti-mouse IgG as the secondary antibody. Detection was done by using ECL(enhanced chemiluminescence) method according to the manufacturer’s protocol [BBI BIOTECH (Shanghai) Co. Ltd, Shanghai, China].

### Colloidal gold labeling

After incubation with S-tagged keratin labeled with immunogold, 1.5 mL cell culture was washed twice and collected at 5000 rpm for 5 min. 1.5 ml of electron microscope fixative solution (3% paraformaldehyde fixative solution, 1% glutaraldehyde) was then added to fix the cells at 4 °C. After 4 h, cell ultrathin sections (50-70 nM) were prepared by Shandon Cryotome E (Thermo Scientific, UN). Ultrathin sections were mounted on 200-300 mesh copper grids covered by 20 uL of 1% H_2_O_2_ for 10 min. The ultrathin sections were washed three times with ddH_2_O for 10 min per wash and incubated with block solution for 15 min at room temperature. The samples were subsequently rinsed with PBS for 5 min to remove the blocking solution. The mouse anti-S-tag monoclonal antibody (1:100 dilution) solution was then added to cover the whole bacterial ultrathin sections for 1 h at room temperature, and further incubated at 4°C overnight. After extensive rinse with PBS for 10 times and 10 min for each, the bacterial ultrathin sections were incubated with 1% BSA solution (pH 8.2) in PBS for 5 min. Colloidal gold-labeled rabbit anti-mouse polyclonal antibody (1:200 dilution) was added dropwise and incubated at room temperature for 30 min. Finally, the sections were rinsed with PBS for 10 times (10 min/rinse), followed by a 5-min wash with 1% lead acetate. Sections were ready for transmission electron microscopy analysis after drying.

### Fluorescence microscope and Transmission electron microscope

After incubation with FITC labeled keratin, cells were washed twice and resuspended in equal volume of IB. Cells were dyed by crystal purple under dark condition prior to fluorescence microscope observation performed with a JEM-2100 transmission electron microscope (TEM) (JEOL Ltd., Japan).

### Keratinase enzyme activity assay

The keratinase enzyme activity was analyzed with the following procedure (11). The reaction mixture contained 1 mL of supernatant from *S. maltophilia* DHHJ culture and 2 mL of 10 mg feather powder in 0.05 mol/L Tris–HCl (pH 7.8) buffer. The mixture was incubated at 40°C for 1 h, and the reaction was stopped by adding trichloracetic acid (100 g/L) to a final concentration of 40 g/L. After centrifugation at 10,000 rpm at 4°C for 10 min, the absorbance of supernatant was determined at 280 nm. One unit of enzyme activity was defined as a change of 0.1 in absorbance at 280 nm.

## Results

### FKM is a major product in feather media culture of *S. maltophilia*

The mechanism underlying the bacterium *S. maltophilia*-mediated feather biodegradation remains undetermined. This led us to trace the intermediates during the feather degradation. We conducted an incubation of *S. maltophilia* bacterial cells with feather media in a 3-day duration and then profiled the protein components in culture media at different time points by SDS-PAGE. Remarkably, a band about 10 kDa was overrepresented in the feather-culture media after the incubation with *S. maltophilia* (Fig. 1b-f), which was not shown in the basic culture media only containing beef extract-peptone (Fig. 1g), indicating a product derived from feather degradation. The protein was recovered from the band through gel-excision then analyzed by mass spectrometry for determining the molecular mass. Mass spectrometry data showed that the target protein has a molecular weight of 10,104 dalton, exactly same as the known keratin monomer.

**Fig. 1.**
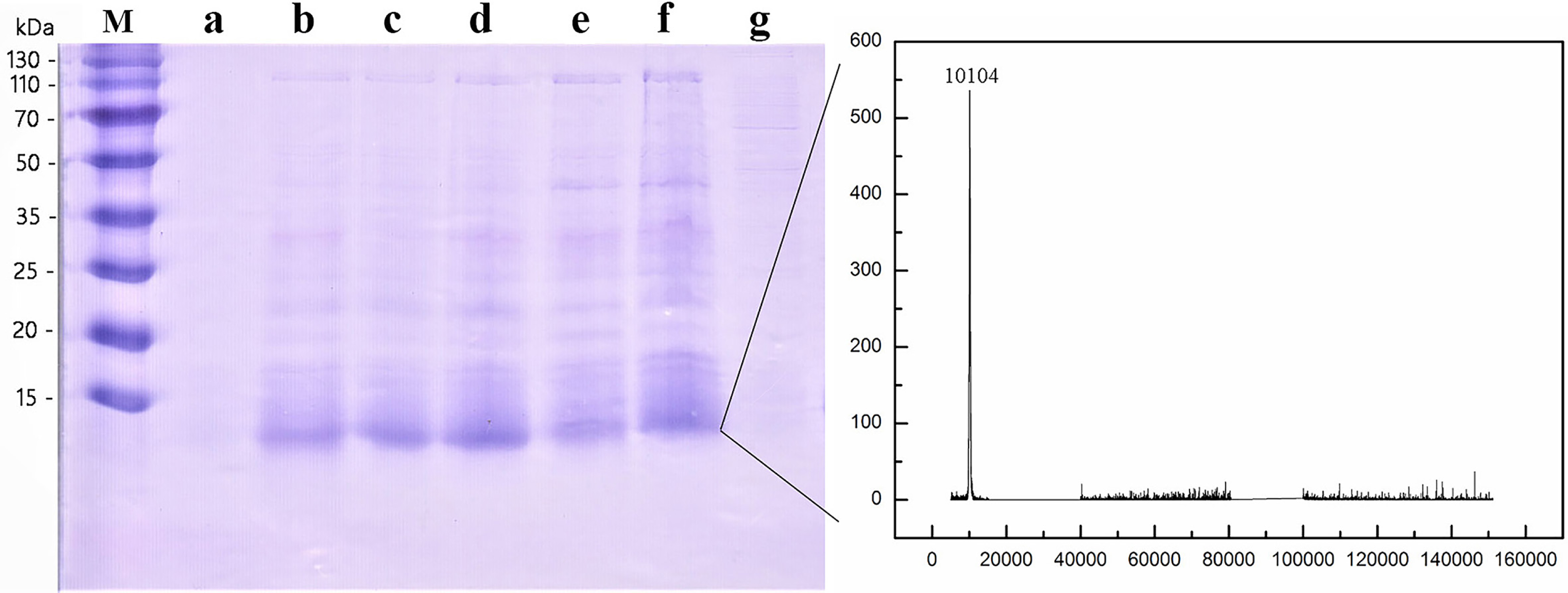
SDS-PAGE detection for the proteins in culture media of *S. maltophilia* DHHJ. a-f. Supernatants of *S. maltophilia* DHHJ cultured with feather media for 0 h, 24 h, 36 h, 48 h, 60 h and 72 h. g. Supernatant of *S. maltophilia* DHHJ cultured with beef extract-peptone media for 48 h.

The band indicated by the arrow was cut and resolved by mass spectrometry for molecular weight and sequence analysis.

### FKMs prepared by chemical degradation or *E. coli* expression and their effect on keratinase induction

To resolve the role of FKM in feather degradation, we prepared FKM with reducing reagent and *E. coli* expression, respectively. To chemically extract FKM from feathers, sodium disulfite (Na_2_S_2_O_5_) was utilized as a reducing reagent for sulfitolysis. In this reaction, the disulfide bonds of keratin were effectively converted into the S-sulfonate anion. The samples treated with reducing agent produced a single abundant band in SDS-PAGE (Fig.2a). This band has a molecular weight of 10,104 dalton resolved by mass spectrometry (MS) (Fig.1), consistent with the theoretical value of feather keratin from *Gallus gallus*.

In order to further explore the action of FKM in feather hydrolysis. FKM was prepared through heterogenous expression in *E. coli*. In the expression vector (Fig.1S), TrxA tag was used for improvement of the protein solubility and His tag was included to facilitate protein purification (Fig.2b). Both the small tags were eventually removed from the final recombinant protein, S-tagged keratin, after cleavage with thrombin protease (Fig.2b).

We examined keratinase activity in *S. maltophilia* cultured in the media with different nitrogen sources. There was a sharp elevation in enzymatic activity in feather meal culture group with maximum induction at 48 h after incubation, followed by a falling phase thereafter, probably due to a feed-back inhibition. We found that FKMs from both chemically-preparation and *E. coli* expression had a similar effect on keratinase activity in the feather-meal samples (Fig. 2c). When cultured with protease K-pretreated feather, the *S. maltophilia* keratinase activity was slightly influenced during culture, revealing a negligible alternation compared to the other treatments. These results suggest that the intact keratin monomer is necessary for stimulation of keratinase activity. In this condition, a modest increase in enzyme activity appeared at 36 hours after culture, probably due to the small quantity of residual keratins from incomplete hydrolysis.

**Fig. 2.**
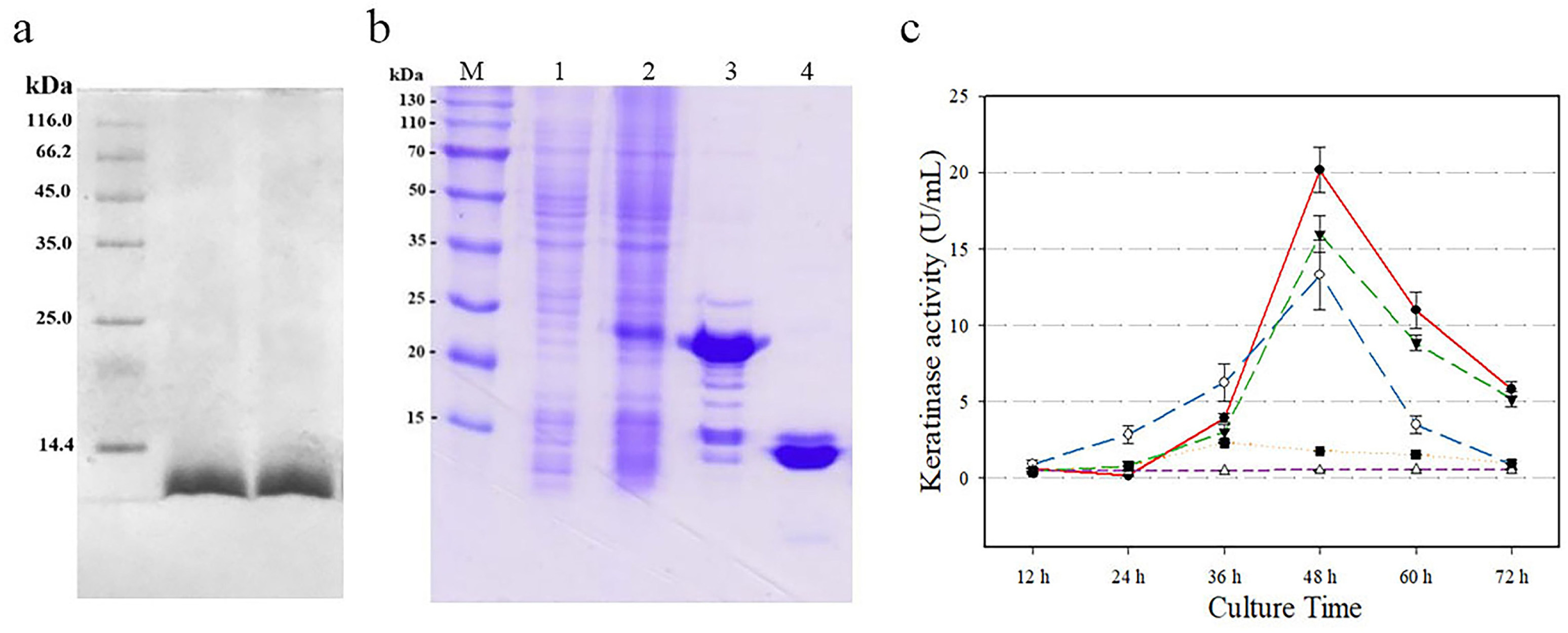
Preparation of artificial FKMs by chemical degradation and heterogenous expression and the effect of the different FKMs on keratinase activity a. SDS-PAGE of FKM made from feathers with chemical solution (2M urea, 0.125 M sodium metabisulfite, 0.05 M SDS, at 60℃ for 30 min). b. SDS-PAGE of the recombinant FKM expressed in E. coli BL21 (a. Without IPTG, b. IPTG induction, c. Purified FKM with His tag before thrombin cleavage, d. Purified S-tagged keratin after thrombin cleavage. c. keratinase activity in S. maltophilia cultured in the media with different nitrogen sources. Feather powder (●); heterogenous expression keratin monomer (○); keratin monomer from feather (▼); keratin monomer degraded by protease K (▪); without inducer (BPD media as a control) (△)

### FITC-FKM binding and internalization by *S. maltophilia* and concurrent increase in keratinase activity

To substantiate cellular entry of FKM, we conducted fluorescence experiment, western blot and immune gold analyses. FITC, a green fluorescence probe, is an ideal tool for tracing dynamic distribution of proteins in cells. We incubated *S. maltophilia* cells with FITC conjugated keratins and visualized fluorescent signal in the treated cells under a fluorescence microscope. Clearly, we observed a large number of cells with green signal in FITC-keratin group but not in control group incubated with FITC alone, suggesting an efficient uptake of FITC-keratin (Fig. 3). This process was apparently corelated with the quantity of *S. maltophilia* cells, as demonstrated by that fluorescence values was correlated with cell density in the culture (Fig S2a). Under a culture condition at 4℃, a temperature below the minimum growth temperature of *S. maltophilia,* we were not surprised to find a remarkable inhibition of KFM-FITC binding (Fig S1b), revealing the temperature sensitivity of the process.

**Fig. 3.**
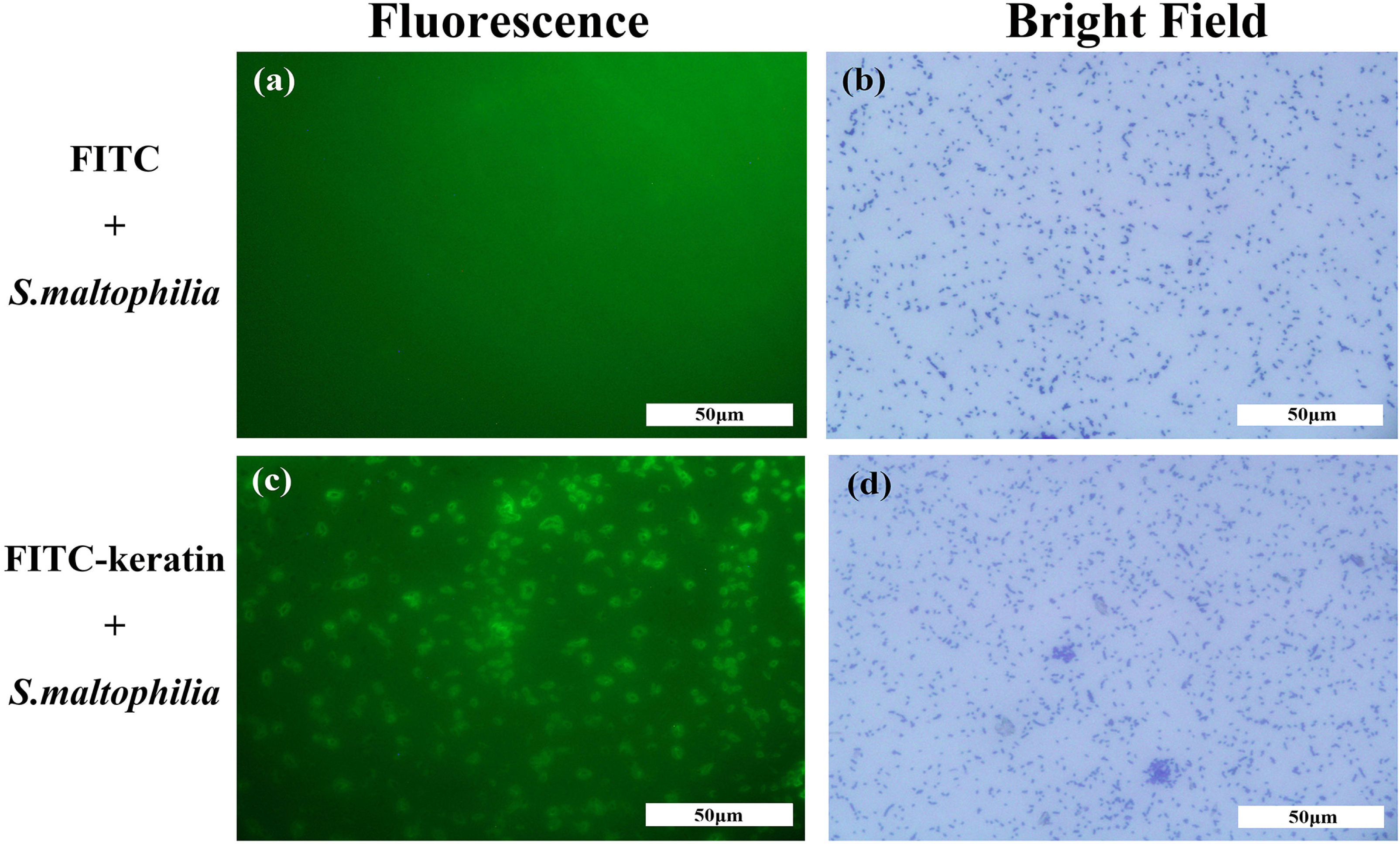
Keratin monomer bind the cells of *S. maltophilia*. Representative images from fluorescence microscope showing *S. maltophilia* incubated with FITC (upper panel, as a control) and keratin-FITC (lower panel)

Further, we cultured *S. maltophilia* with the addition of keratin-FITC and found the specific binding of keratin but not FITC (as a control) by the cells. Fluorescence intensity dramatically increased in the cells after 24-h incubation with keratin-FITC, which remained at a higher level thereafter. Meanwhile, we observed that keratinase activity spiked at 48 h after keratin-FITC incubation (Fig 4). While the *E. coli* group did not show similar results (Fig 4).

**Fig. 4.**
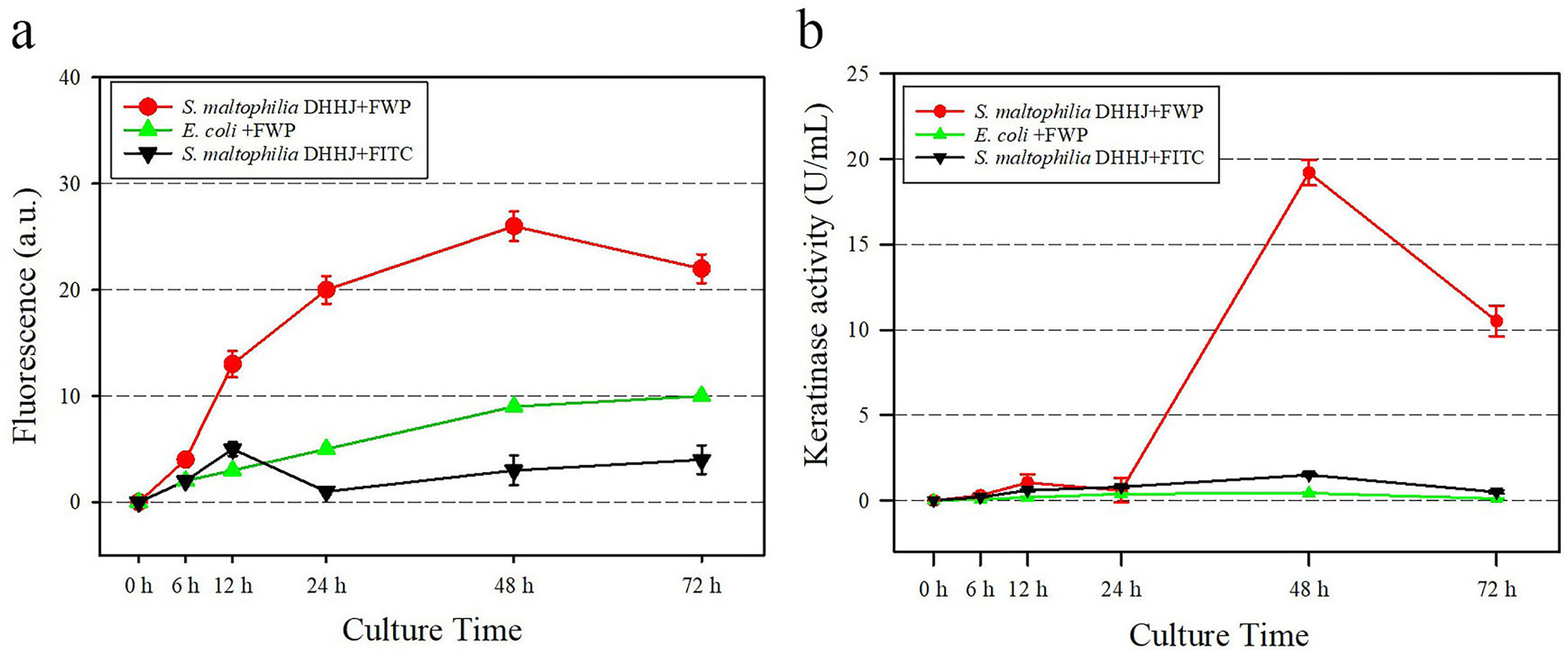
Keratin monomer stimulates keratinase production in *S. maltophilia* through cellular binding. *S. maltophilia* cells specifically bind keratin monomer and consequently stimulate keratinase production. The culture medium of *S. maltophilia* DHHJ was added with keratin-FITC (400 μg/mL, FWP) or FITC alone. *E. coli* was cultured with keratin-FITC (400 μg/mL, FWP) containing medium. (a. fluorescence intensity of the cells; b. keratinase activity)

We next confirmed the cellular entry of FKM by western blot analysis. As shown in Fig. 5a, keratin monomer was detected in whole cell lysate as well as in both membrane and intracellular fractions. The uptake of the S-tagged keratin monomer by *S. maltophilia* cells was confirmed by western blot analysis (Fig. 5a) and immunogold labeling analysis (Fig. 6). Keratin monomer could be identified in total protein, membrane as well as inner cellular components after its incubation with *S. maltophilia* cells. The western blot data suggested that FKMs were translocated into cells by crossing the membrane through an undefined transport mechanism. It seems that *S. maltophilia* cells control FKM internalization in a protein-cell specific mode, because, in control experiments, FKM was not internalized by *E. coli* and similarly a S-tagged control protein was not taken into *S. maltophilia* either (Fig. 5b). Immunogold labeling cryosections of high-pressure frozen cells were used to investigate whether the internalized FKM was associated with intracellular structures. As shown in Fig. 6a and 6b, gold particles, representing presence of FKMs in the cells, were often seen in an aggregating form in either longitudinal or cross sections. We also observed that gold particles did not randomly spread, but tended to gather at certain structures, similar to infolded cytoplasmic membranes (Fig. 6C), probably revealing the initial stage of the FKM internalization process. There were many vesicles on the *S. maltophilia* cell surface during FKM incubation. This may suggest the association between FKM internalization and vesicle function.

**Fig. 5.**
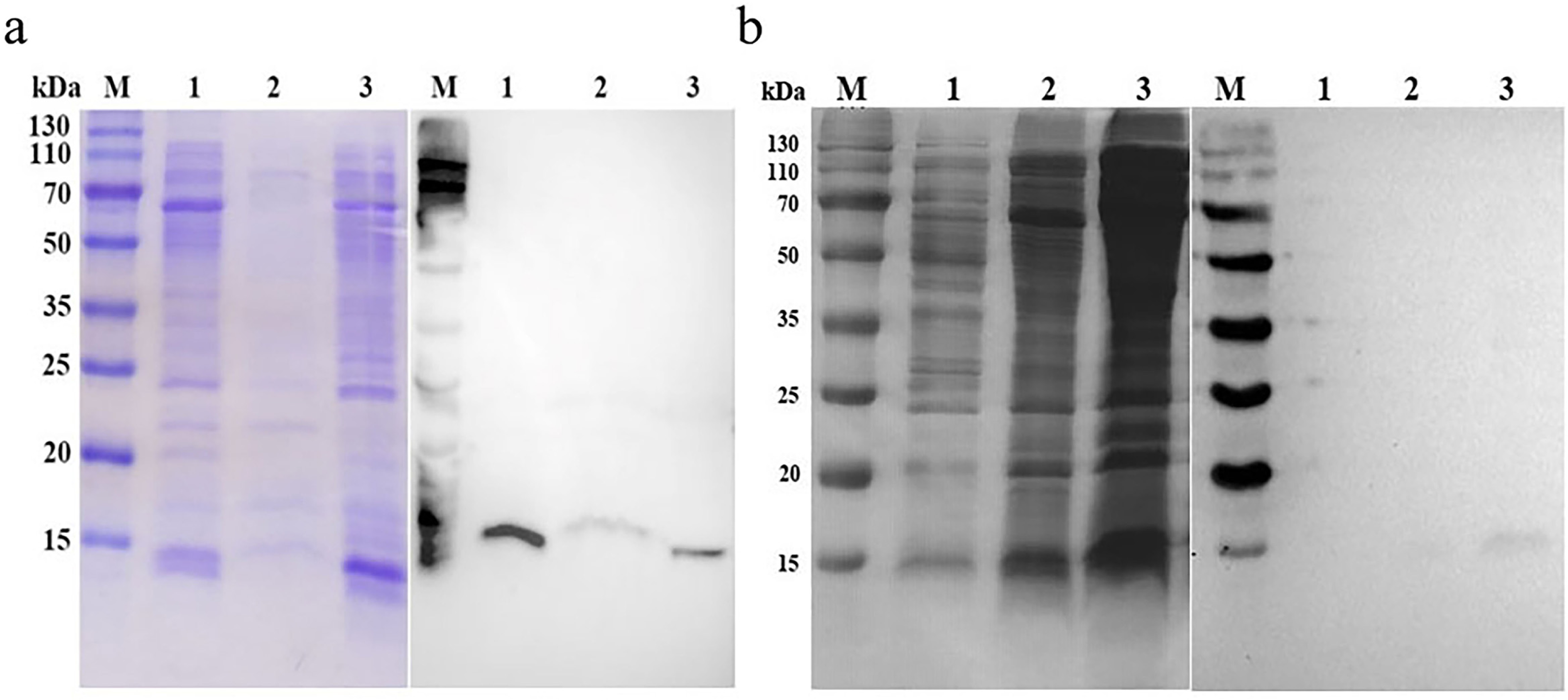
Western blot analysis showing uptake of keratin monomer (∼10 kDa) by *S. maltophilia*. a. Cells were coincubated with keratin monomer for 4h. After washing, cell lysates were separated on an SDS-PAGE gel (left), then blotted and probed with an anti-S tag antibody (right). 1. Total proteins 2. Membrane proteins 3. Intracellular proteins b. *E. coli* (1) and *S. maltophilia* (3) cells were coincubated with keratin monomer for 4h. *S. maltophilia* cells (2) were coincubated with S-tagged Ca-tif1(as a control). Protein separation by SDS-PAGE gel shown on the left and western blot probed with an anti-S tag antibody shown on the right.

**Fig. 6.**
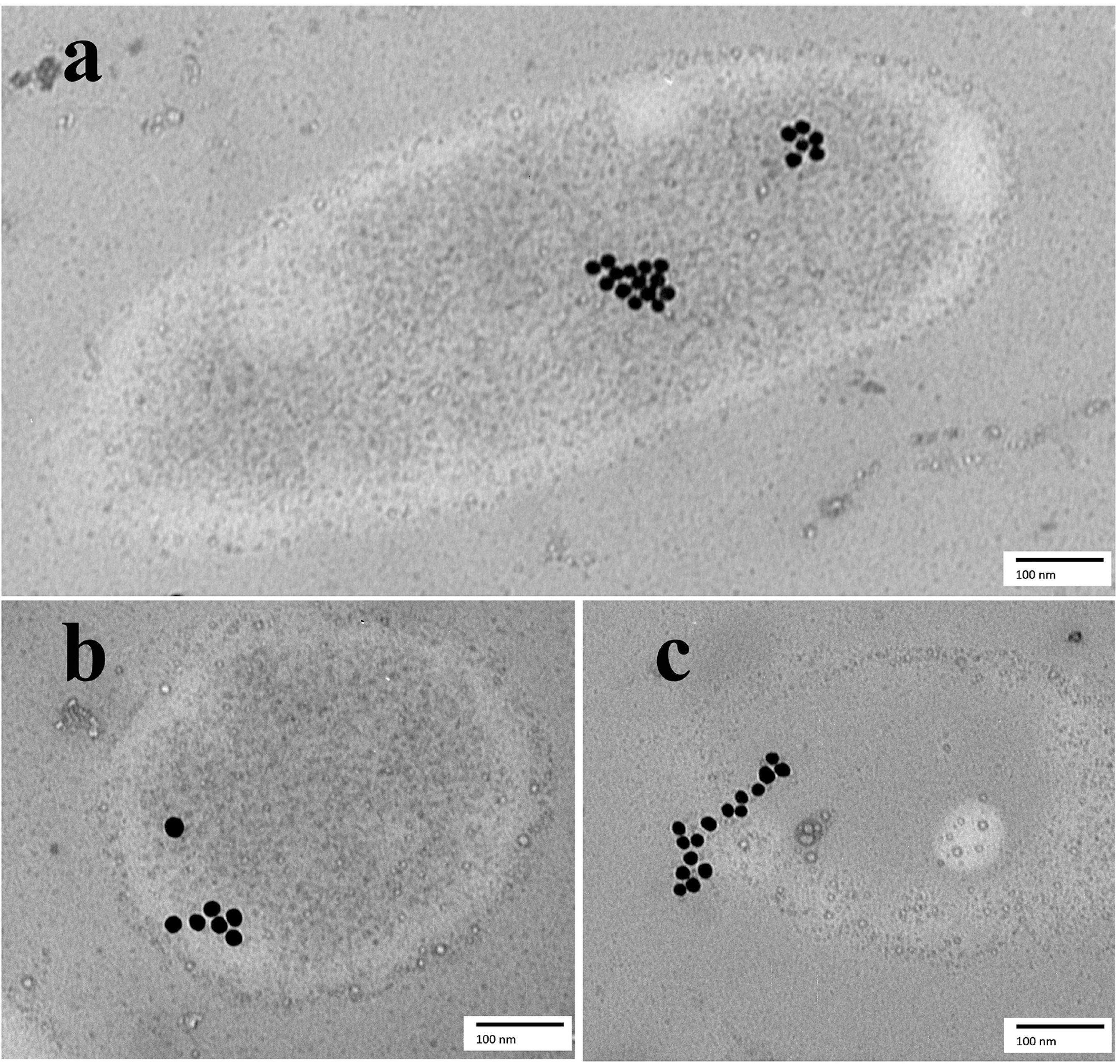
TEM of high-pressure frozen cryosections of *S. maltophilia* cells showing association of keratin monomers with membranes and cytoplasm. (a) Cells were labeled with immunogold to detect keratin monomers, which were seen in the cytoplasm (arrowhead). (b, c) Sections showing infolding of cytoplasmic membrane (arrowhead) and cytoplasm associated with keratin monomer (gold particles). TEM image of a section of high-pressure frozen and cryosubstituted cells of *S. maltophilia*, immunogold assay to detect FKM with S-tag via anti S-tag antibody and secondary antibody conjugated with 10 nm colloidal gold.

Taken together, these results demonstrated that feathers can initially be hydrolyzed to FKM when supplied as the nitrogen source in *S. maltophilia* culture medium. FKM can bind to cells membrane and subsequently enter cells to undergo complete hydrolysis. This hypothesis is also supported by the finding of induction in keratinase activity by FKM. This process is energy dependent and BSA greatly diminished uptake of FKM, while nucleic acid did not affect FKM uptake (data not shown).

## Discussion

Although accumulating evidence have showed the efficacy in feather keratin breakdown by using keratinases-producing microogamisms(10), these technologies may face challenge in scaling up for industrial application. Keratin fermentation also needs special equipment, prolonged reaction time, and high energy input for maintenance of agitation and reaction temperature. More importantly, considerable amount of end products will be eventually turned into microbial metabolites in keratin fermentation, obviously contrary to the recycling purpose. To circumvent these disadvantages, direct application of keratinases is considered a better substitution. However, a thorough understanding of the mechanism underlying keratinase-mediated feather degradation is the prerequisite for establishing enzyme-based biodegradation in practical applications. To this end, we conducted the mechanistic study by using the S. *maltophilia*-feather meal medium model. We found a 10 kDa water-soluble protein as a major hydrolytic product derived from feathers in *S. maltophilia* culture medium, which was further confirmed to be the keratin monomer. Multiple lines of evidence from the experiments based on both chemically generated and bacterially expressed keratin monomers indicate that the complete hydrolysis of keratin requires the internalization of keratin monomers into *S. maltophilia* cells.

The detailed mechanism underlying the formation of extracellular keratin monomer by *S. maltophilia* remains elusive. This process may need the involvement of disulfide reductases, keratinases, and other peptidases. Microbial keratinases can simply be classified as serine- or metallo-type proteases, whereas these keratinolytic enzymes actually have shown numerous characteristics in terms of biochemical properties. The molecular masses of the known keratinases vary in a wild range. For example, *Streptomyces albidoflavus* produces an 18 kDa keratinases (22), while the alkaline extracellular keratinase secreted by *Kocuria rosea* has a molecular mass of >200 kDa (23, 24). The optimal pH and temperature for keratinase activity from different sources show a great discrepancy (2). The microbial keratinases include extracellular, cell-bound, and intracellular enzymes (25). Some keratinases are monomeric while others can be a dimer (2). These dissimilarities in biochemical features expand the complexity of proteolytic mechanisms of keratinases. Some keratinases were reported to be able to hydrolyze keratin by themselves, whereas more studies found solo keratinases are not sufficient for digestion of keratinous substrates and cooperation with other components is necessary. Obviously, the mechanism proposed in this study is in favor of the notion about involvement of multiple factors in keratolytic process.

Currently little is known about the pathway through which *S. maltophilia* cells assimilate keratin monomer for further degradation. The transport of peptides in microorganisms is an important physiological process involved in nutrient assimilation and cell-cell communication (26). Oligopeptide permease (OPP) systems have been extensively studied in microbial peptide uptake. Lactic acid bacteria (LAB), the major microorganisms for milk fermentation, are a well-studied model for bacterial peptide transportation (27). LAB acquire essential amino acids from milk during their growth by employing extracellular proteolytic enzymes to hydrolyze casein and intrinsic transporting machineries to assimilate the degraded peptides (28), which depend on the function of three systems: Opp, DtpT, and Dpp. The LAB Opp systems transport complex peptides containing 5 to 20 or more residues (29). The other transporters DtpT and Dpp in LAB mainly deal with smaller peptides such as di-, tri- or tetrapeptides (29). In this study, feather keratin monomer was found to be assimilated into *S. maltophilia* cells, which has a molecular weight of 10 kDa, likely far beyond the capability of the ABC transporter system, suggesting that other types of transportation pathway for large peptides/proteins may exist in *S. maltophilia*.

Studies have shown that microorganisms adopt other mechanisms for macromolecule transport. For example*, Gemmata obscuriglobus* was reported to uptake GFP (238-aa), much larger than KFM, through an energy-dependent endocytosis-like process, in which internalized proteins were wrapped by vesicle membranes (30). Here the data from scanning electron microscopy also showed the gold particles (S-tagged keratins) associated with the infolding membrane structure and the uptake process was indispensable to the bacterial growth, as indicated in a low temperature inhibition assay. Although peptide internalization in *S. maltophilia* shares some similarities to that of *Gemmata obscuriglobus,* more experiments need to be done to elucidate the exact mechanism.

In summary, this study provided new insights in the coarse-to-fine process of feather biodegradation in *S. maltophilia* DHHJ. We postulated that the whole degradation pathway is precisely controlled and well organized, which is compartmentalized as extracellular and intracellular events. The reaction starts from extracellular conversion of insoluble feathers to soluble keratin monomer, in which enzymes catalyzing reduction of disulfide bonds are likely recruited at initial stage and some proteinases may also be participated in the action, which remain to be characterized. We hypothesize that the biological meaning of the initial step in feather degradation in *S. maltophilia* is to convert insoluble substrate to soluble keratin monomers, which are possible to be relocated into the bacterial cells, where the keratin monomers can be further broken down. So far, we do not know whether this mode of feather degradation is only restricted to *S. maltophilia* and whether *S. maltophilia* cells can degrade α-keratin in the same way. All these questions are worthy of further investigation.

## Acknowledgments

This work was supported by the National Natural Science Foundation of China under 31570106 and 31000989.

---

**Fig. S1.**
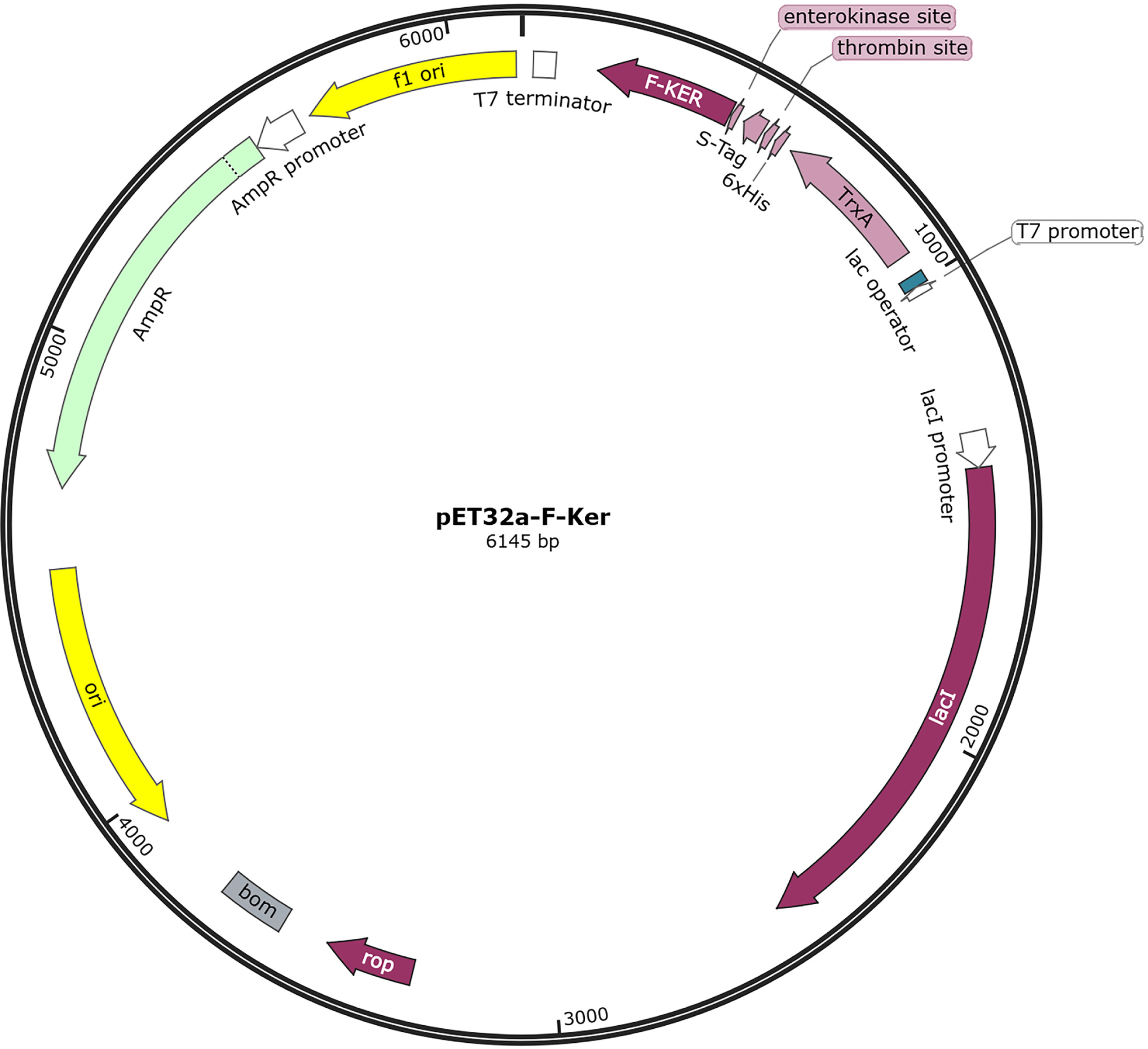
Schematic diagram of pET-32(+) with feather keratin gene (Gene ID: 769269)

**Fig. S2.**
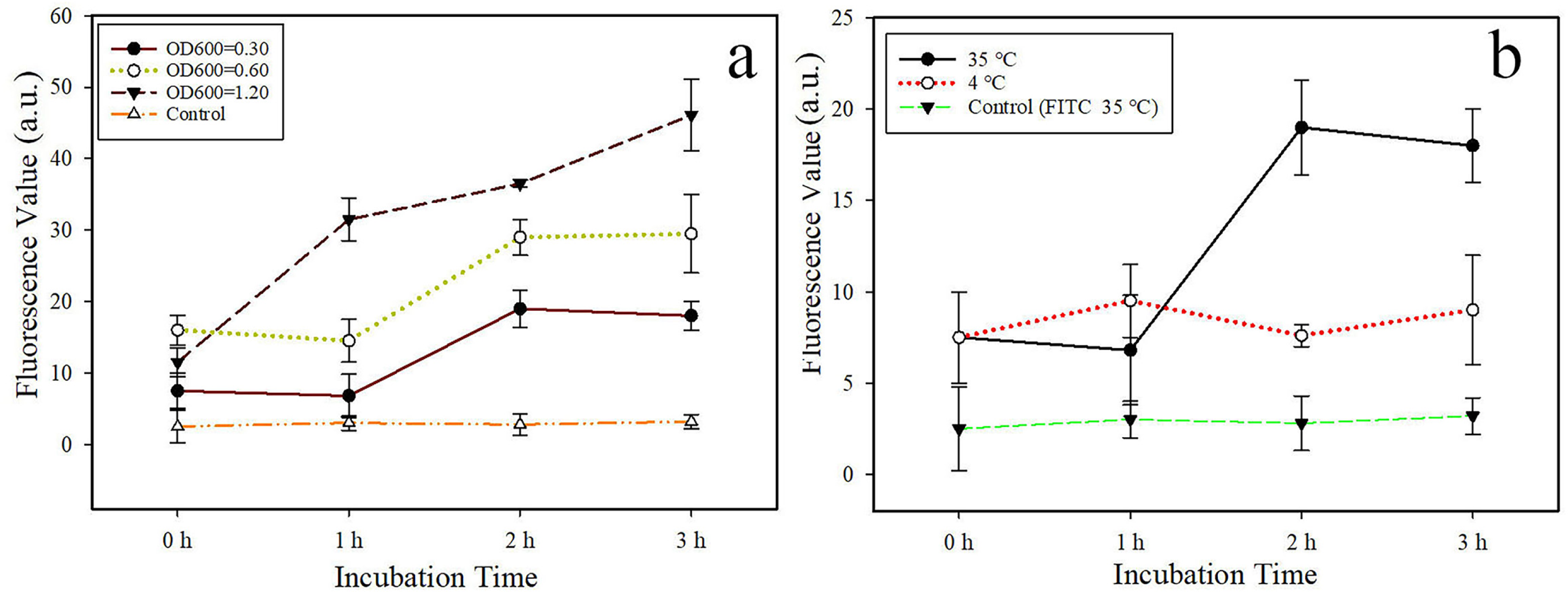
Effect of cell density and temperature on cellular binding of keratin monomer to *S. maltophilia.* a. Keratin monomer is binding with different cells concentration. b. Keratin monomer is binding with cells at different temperatures.

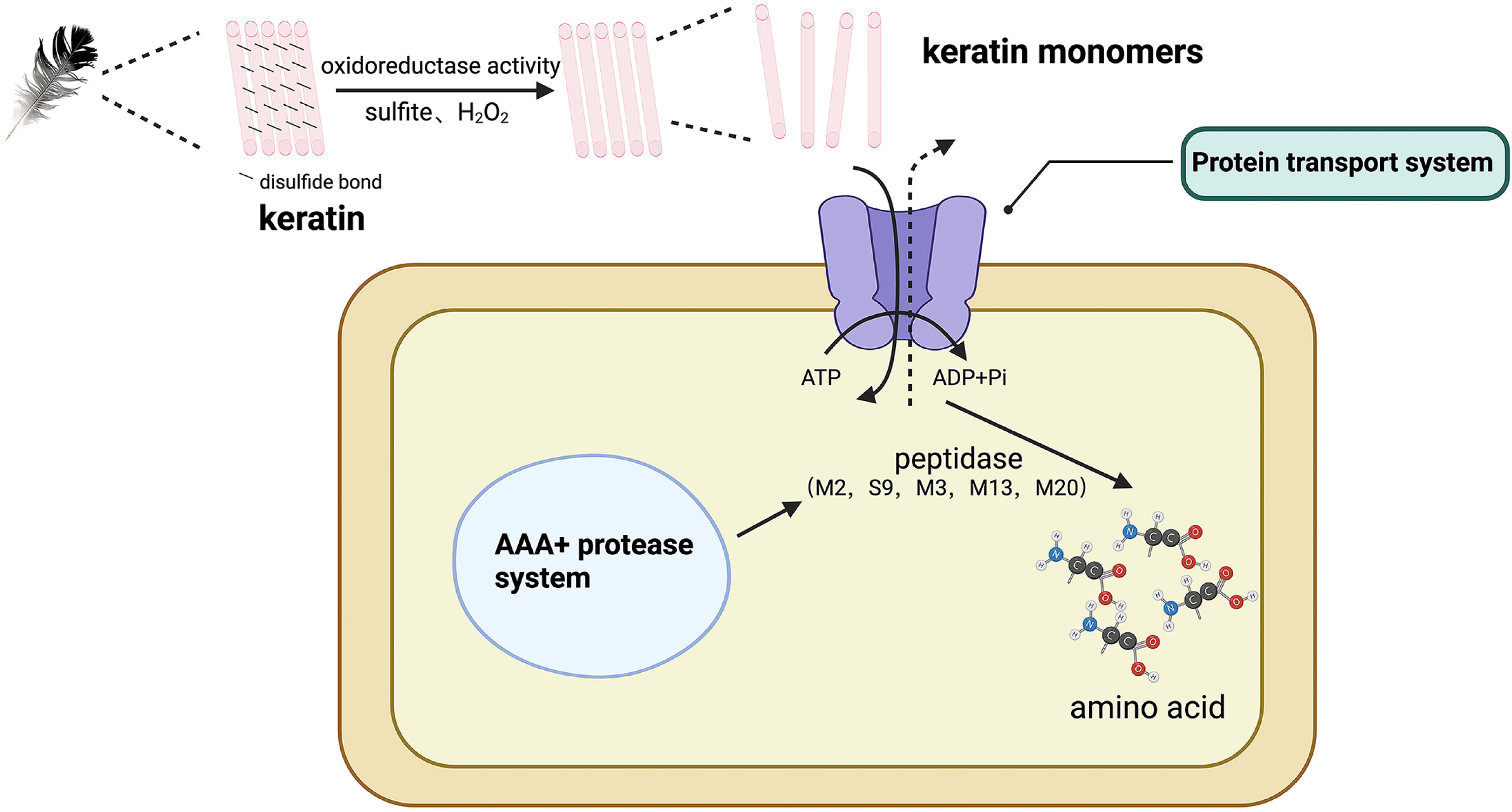

